# Long-term independent use of an intracortical brain-computer interface for speech and cursor control

**DOI:** 10.1101/2025.06.26.661591

**Authors:** Nicholas S. Card, Tyler Singer-Clark, Hamza Peracha, Carrina Iacobacci, Xianda Hou, Maitreyee Wairagkar, Zachery Fogg, Elena Offenberg, Leigh R. Hochberg, David M. Brandman, Sergey D. Stavisky

**Affiliations:** University of California, Davis, Dept. of Neurological Surgery, Davis, CA, USA; Dept. of Neurology and Neurosurgery, University Medical Center Utrecht Brain Center, Utrecht University, Utrecht, The Netherlands; Sch. of Engin. and Carney Inst. for Brain Sci., Brown Univ., Providence, RI, USA; VA RR&D Ctr. for Neurorestoration and Neurotechnology, Providence VA Med. Ctr., Providence, RI, USA; Dept. of Neurol., Massachusetts Gen. Hosp., Boston, MA, USA

**Author notes:** Co-senior authors (David M. Brandman, Sergey D. Stavisky).

## Abstract

Brain-computer interfaces (BCIs) can provide naturalistic communication and digital access to people with severe paralysis by decoding neural activity associated with attempted speech and movement. Recent work has demonstrated highly accurate intracortical BCIs for speech and cursor control, but two critical capabilities needed for practical viability were unmet: independent at-home operation without researcher assistance, and reliable long-term performance supporting accurate speech and cursor decoding. Here, we demonstrate the independent and near-daily use of a multimodal BCI with novel brain-to-text speech and computer cursor decoders by a man with paralysis and severe dysarthria due to amyotrophic lateral sclerosis (ALS). Over nearly two years, the participant used the BCI for more than 3,800 cumulative hours to maintain rich interpersonal communication with his family and friends, independently control his personal computer, and sustain full-time employment – despite being paralyzed. He communicated 183,060 sentences – totaling 1,960,163 words – at an average rate of 56.1 words per minute. He labeled 92.3% of sentences as being decoded at least mostly correctly. In formal quantifications of performance where he was asked to say words presented on a screen, attempted speech was consistently decoded with over 99% word accuracy (125,000 word vocabulary). The participant also used the speech BCI as keyboard input and the cursor BCI as mouse input to control his personal computer, enabling him to send text messages, emails, and to browse the internet. These results demonstrate that intracortical BCIs have the potential to support independent use in the home, marking a critical step toward practical assistive technology for people with severe motor impairment.

## Introduction

Loss of the ability to speak and use digital devices profoundly impacts the independence and quality of life of people with severe motor impairments caused by conditions such as amyotrophic lateral sclerosis (ALS) or brainstem stroke^1–3^. Although a variety of augmentative and assistive communication (AAC) technologies exist to support communication and computer use for people with paralysis, these tools are often slow, fatiguing, unreliable, or require frequent intervention from carepartners trained in their use^4^. Recent progress in brain-computer interfaces (BCIs) has shown a promising alternative pathway toward restoring naturalistic communication and digital access, by decoding neural activity during attempted speech or movement into words or digital actions^5–13^.

Intracortical BCIs, which record neural signals at the resolution of action potentials, have enabled the highest-performance decoding to date of cursor^9^, handwriting^6^, typing^13^, and speech^5,12^. Intracortical BCIs have traditionally focused on decoding either speech or hand movement-based control, often relying on separate cortical regions and independent computational architectures. Recent studies have shown that speech-related activity in the motor speech cortex can be decoded into text with high speed and accuracy, enabling brain-to-text communication at near-conversational rates^5,7,10^. Other work has demonstrated that neural signals related to intended arm and hand movements can drive high-performance cursor control^9,14–17^, and one recent study has shown that both capabilities can be decoded from the speech motor cortex^18^. However, most prior intracortical BCIs have addressed speech or cursor control in isolation, required frequent recalibration to maintain accuracy, and/or required researcher oversight to don and doff the system. Moreover, existing BCIs have typically been evaluated in short-term research settings, without demonstration of extended high performance over many months of independent use.

These limitations highlight two major barriers to the clinical translation of intracortical BCI technology. First, the transition from a scientific demonstration to a practical communication device requires users (and their care partners) to be able to operate the system independently in their homes, without daily support from researchers or technicians. Second, neural recording sensors and decoding performance must remain stable over long durations despite the neural signal variability associated with chronic implants^19–21^. Ideally, little to no time should be spent asking the user to interrupt their use to recalibrate the decoder.

Here, we report the long-term, independent use of a multimodal intracortical BCI that enables both brain-to-text speech and computer cursor control. With care partner assistance largely restricted to donning and doffing the hardware and initiating the software, a man with paralysis and severe dysarthria due to ALS used the system in his home nearly every day for 19 months, accumulating over 3,800 hours of independent use. The system incorporated novel decoding architectures for both speech and cursor control, as well as multiple software features that enabled independent use and continuous decoder finetuning. For speech, we developed a transformer-based brain-to-text decoder that outperformed prior RNN-based models, requiring little-to-no daily calibration and achieving a new state-of-the-art word accuracy of 99.2% in a prompted word Copy Task (125,000 word vocabulary). For cursor control, we implemented an RNN-based decoder that matched or exceeded the performance of linear models, also with reduced calibration requirements. Using this system, the participant communicated more than 180,000 sentences at conversational speeds and used the speech and cursor decoders together to independently operate his personal computer – sending messages, browsing the internet, participating in video calls, and maintaining full-time employment – despite being paralyzed.

These results demonstrate that intracortical BCI systems, equipped with advanced decoding algorithms, can now support rich, independent digital and in-person communication in real-world settings. These results are a significant step toward delivering a practical assistive technology for people with severe speech and motor impairments.

## Results

### Overview

A 45-year-old man with paralysis and severe dysarthria due to ALS (‘T15’) enrolled in the BrainGate2 clinical trial (ClinicalTrials.gov number, NCT00912041) in 2023. Four microelectrode arrays (64 electrodes each) were surgically placed in T15’s ventral precentral gyrus (speech motor cortex) to record intracortical neural activity via percutaneous wired connections (Fig. 1A, Fig. 1B, left). Neural signals (Fig. 1B, center) were processed by three parallel neural decoders for real-time translation of T15’s neural activity into intended words, cursor movements, and click gestures (Fig. 1B, right). A continuously running “brain-to-text” decoder detected and decoded attempted speech into the most likely word sequences. Two additional decoders – activated by a gaze-controlled toggle – translated right-hand motor imagery into two-dimensional cursor control and discrete clicks. Details of decoder architectures and training procedures are provided in the Methods.

**Figure 1.**
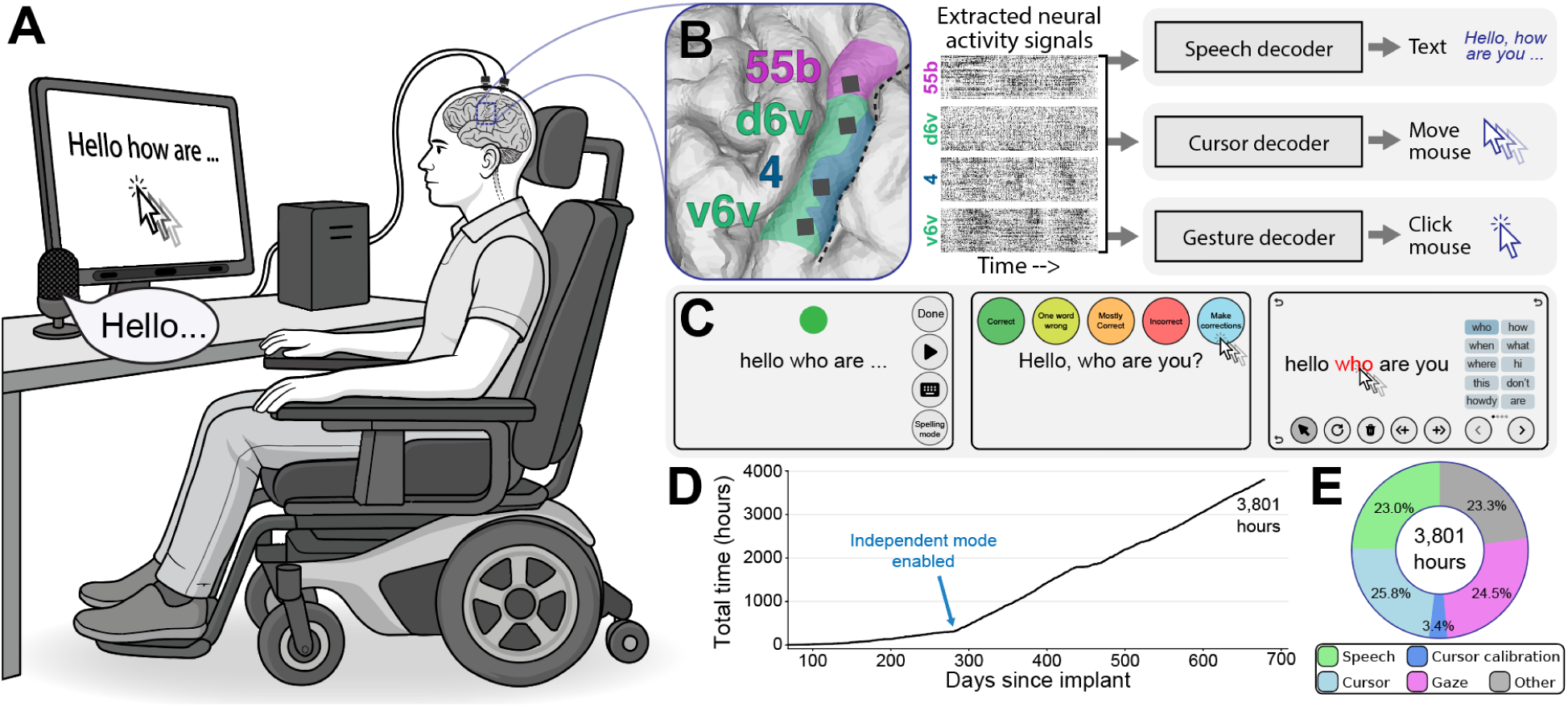
Independent use of the multimodal intracortical BCI. **(A)** Schematic of the participant using the multimodal intracortical BCI to control their personal computer. As the participant tried to speak, his neural activity was decoded into words on a screen. He could also control his computer by attempting to move or squeeze his hand. The system also integrated eye tracking to enable the participant to select on-screen buttons by looking at them for a short time. **(B)** Locations of four 64-microelectrode arrays in the participant’s dominant ventral precentral gyrus (*left*). The dotted line is the dominant central sulcus. Each electrode records activity from individual neurons and local field potentials. Recorded neural signals were processed into binned sequences of neural features (*middle*) and inputted to speech, cursor, or gesture decoders to decode the user’s intended words, cursor movements, or clicks (*right*). **(C)** User interface examples during speech decoding (*left*), sentence rating (*middle*), and sentence correction (*right*). **(D)** Cumulative hours of system usage over time. Independent use began at post-implant day 281. **(E)** Time distribution of how the system was used across 3,801 hours sourced from all personal use sessions.

The BCI system in T15’s home was a set of networked research computers^22^ mounted on a mobile cart, located in his bedroom or living room. T15’s care partners were trained to safely connect the neural recording equipment and power on the computer system, which was then automatically initialized through custom software. This allowed T15 to operate the BCI without assistance from any member of the research team. Daily setup took approximately 20 minutes, after which T15 used the system continuously for up to 19 hours without additional assistance. Once initialized, T15 operated the system independently using neural decoding, supplemented by eye gaze tracking to facilitate user-interface control. The speech decoder software ran continuously in the background and displayed decoded words on-screen in real time (Fig. 1C, left). After each utterance, T15 was given the option to make corrections to the decoded text through a custom user interface via gaze or cursor input (Fig. 1C, right). He then rated decoding accuracy with the following options: “correct,” “one word wrong,” “mostly correct,” or “incorrect” (Fig. 1C, middle).

We previously quantified T15’s natural speech and effective communication rates with his preferred AAC device^5^. Rather than relying on his existing communication strategies, T15 chose the BCI as his primary mode of communication and computer interface. He used it to converse with family, friends, colleagues, and clinicians – in person, over video calls, and through digital tools such as email and text messaging. He also used it for both professional and recreational computer access. Before post-implant day 281, usage was limited to 2-4 weekly sessions requiring a research assistant (author C.I.) to oversee setup and use (3.7 hours/day average). After the investigational device exemption (IDE) was modified allowing his care partners to don and doff the system without the scientific team being present on day 281, T15 and his care partners could initiate system use independently, resulting in a marked increase in usage to 9.5 hours/day average (Fig. 1D). As of 678 days after the implant surgery, he had accumulated over 3,800 hours of BCI use in 444 days (Fig. 1E).

### Speech decoding

After implant day 281, T15 used the brain-to-text decoder to communicate independently on a near-daily basis. The decoding pipeline (Fig. S1) converted neural features into English phoneme probabilities every 80 ms via a neural network (Fig. S2), followed by language modeling to generate the most likely word sequences drawn from a vocabulary of over 125,000 English words. The decoder was continuously recalibrated in the background to compensate for slow shifts in neural activity and maintain high decoding accuracy. The output was shown on-screen after each word and was optionally synthesized at the end of a sentence using a text-to-speech system trained to match T15’s pre-ALS voice. Over the course of the study, we iteratively improved the decoder architecture – starting from the approach described in ^5^ – to increase accuracy, reduce explicit calibration requirements, and improve robustness to signal variability (Fig. S2).

Decoding accuracy was self-rated by T15 after each sentence and remained relatively stable over months (Fig. 2A, top). The ability to make corrections using the custom-built graphical user interface, controlled via gaze or cursor, became available at post-implant day 227. We observed that sentence accuracy was influenced by multiple factors, including fatigue (Fig. S5), attempted speaking rate (Fig. 2A, subplot 2), sentence length (Fig. 2A, subplot 3), and topic. System updates – including new feature implementations, bug fixes, and decoder upgrades – also contributed to performance variability (Fig. S2). Across 183,060 sentences, 53.3% were decoded completely correctly, 12.9% were corrected by the participant, and 26.1% were mostly correct (Fig. 2B). The overall trend of decoding performance improved over time, with later months showing a higher proportion of correct sentences (rank sum test; p<0.001; Fig. 2C) and increased sentence length (p<0.001; Fig. 2A, subplot 3). We note that T15 often strung multiple sentences together at a time into a single utterance – the longest correctly decoded utterance was 215 words (Fig. S4) – which had the effect of lowering the self-rated accuracies (as an utterance gets longer, the probability of it being decoded 100% correct decreases).

**Figure 2.**
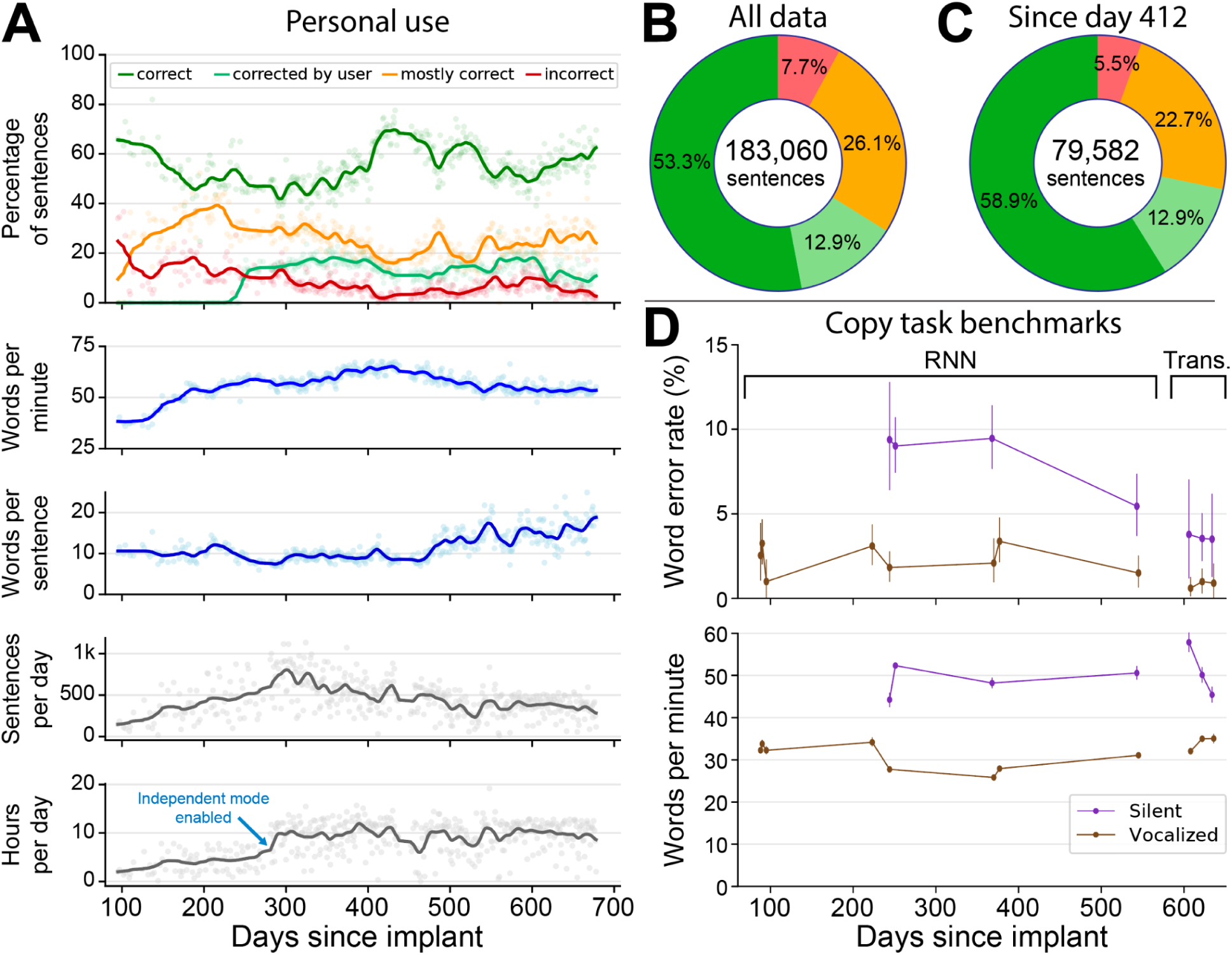
Speech decoding. **(A)** “Personal use” speech decoding usage and accuracy statistics by day. The first subplot shows the percentage of sentences that the participant reported to be (i) immediately correct, (ii) initially not correct but corrected through the user interface, (iii) mostly correct, or (iv) incorrect. Subplots 2-5 show the daily average rate of communication (words per minute), average words per sentence, total sentences communicated, and total hours of system use. **(B)** Distribution of self-reported sentence correctness among 183,060 sentences sourced from all personal use sessions. Color legend is the same as the top plot in panel **(A)**. All self-reported sentence accuracy ratings were made at the participant’s discretion. **(C)** Self-reported sentence accuracy ratings limited to the 238 most recent days of usage (post-implant days 412 to 678; 79,582 sentences). Day 412 was chosen as the cutoff point because of a substantive update to the continuous decoder finetuning algorithm. **(D)** Word error rate (*top*) and speaking rate (*bottom*) in periodic Copy Task benchmark sessions using silent (*violet*) and vocalized (*brown*) speaking strategies. The final 3 benchmark sessions used the transformer-based decoder, all prior sessions used the RNN-based decoder.

We did not instruct T15 to use a particular speaking strategy during personal use of the system; instead, he was encouraged to use whichever approach felt most sustainable, natural, and effective. The opportunity to develop a strategy that minimizes user fatigue was especially important, given the known challenges of fatigue in patients with late-stage ALS. Initially, he employed attempted vocalized speech, producing sound while speaking. Over time, he transitioned to a “silent speech” strategy – making facial muscle movements without phonating. This strategy change led to increased speaking speed, from ∼30 to over 50 words per minute (WPM; Fig. 2A, subplot 2). To benchmark longitudinal performance of the system, and to quantify the effect of silent vs. overt speaking strategies, we periodically asked him to repeat sentences presented on a screen in a Copy Task (Fig. 2D). During vocalized speech, accuracy exceeded 99% at 30.6 WPM average; with silent speech, accuracy reached 96.5% at 49.7 WPM average. Note that personal use and benchmark sessions after post-implant day 600 utilized a more accurate transformer-based phoneme decoding architecture, while prior sessions used a recurrent neural network-based model (Fig. S2).

### Cursor decoding

Although T15’s microelectrode arrays were placed in the ventral precentral gyrus (i.e., speech motor cortex), we previously described^18^ how he performed two-dimensional cursor movements and click events using motor imageries (hand, or body-part-agnostic) that traditionally are used when decoding point-and-click from dorsal (hand) precentral gyrus. In structured Grid Task assessments, the system achieved up to 3 bits per second of throughput, comparable to performance levels reported with arrays placed in hand motor areas^9,17^.

Building on this finding, we developed a custom software interface linking the BCI system to T15’s personal computer, enabling him to control the mouse pointer with decoded neural signals (Fig. 3A). By enabling him to “copy/paste” the text decoded by the speech BCI, he could use his computer for a range of daily activities including writing, web browsing, and participation in video calls. Cursor control functionality was added on post-implant day 358, after which T15 used the feature for an average of 121.0 ± 66.9 (mean ± standard deviation) minutes per day (Fig. 3B).

**Figure 3.**
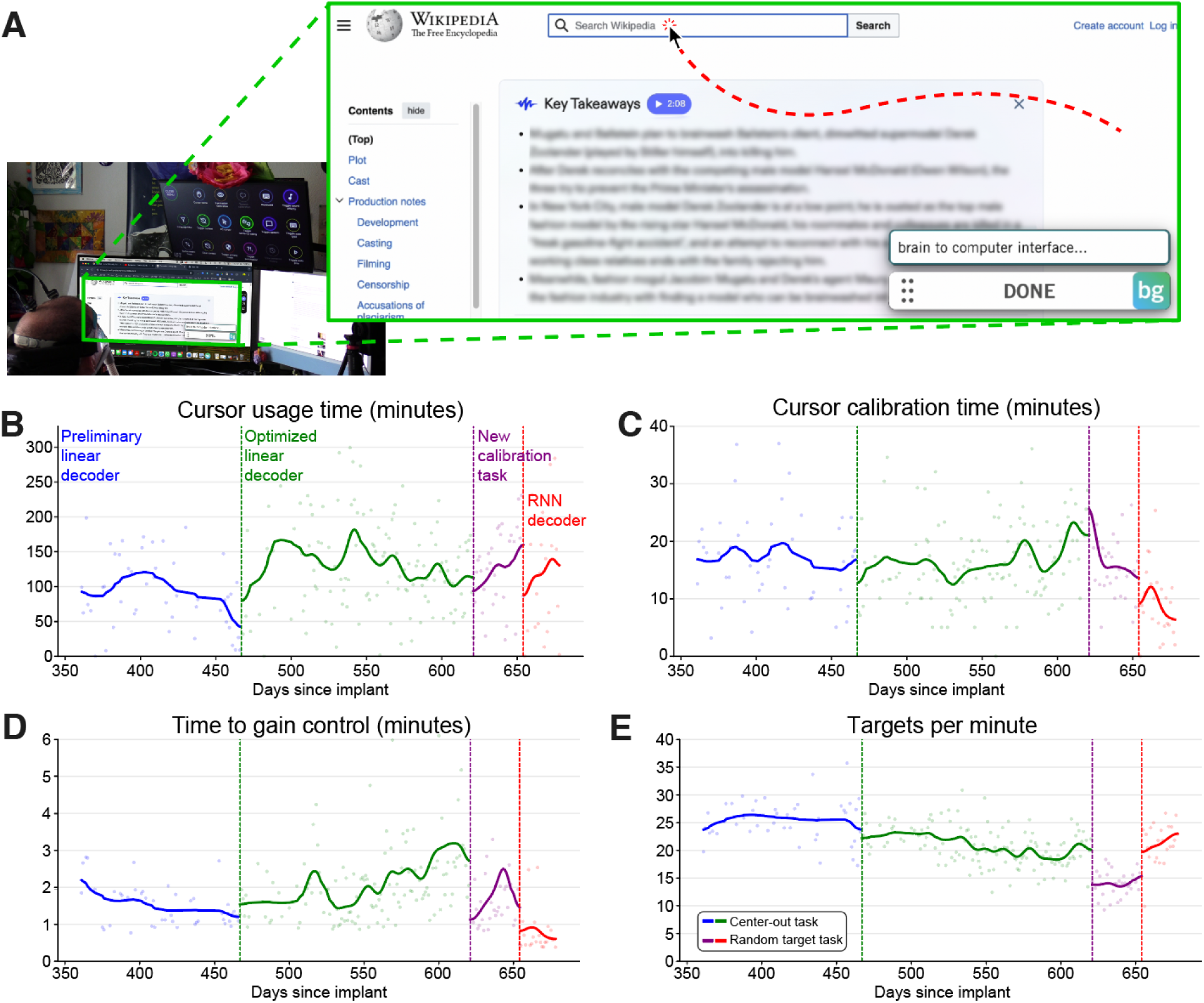
Computer control with the cursor BCI. **(A)** Graphic of T15 using the BCI with his personal computer to search wikipedia for “brain to computer interface”. The current decoded words are shown in the bottom right hand of the screen via our custom BGHome software. T15 can move and click the mouse via the cursor decoder to select the search bar. **(B)** Amount of time T15 used the cursor BCI to operate his computer. The cursor BCI served as T15’s primary method of computer control. Blue and green data points utilized a linear decoder (preliminary or optimized, respectively) and a center-out-and-back calibration task. Purple data points used the optimized linear decoder with a random target calibration task. Red data points used an RNN-based decoder with a random target calibration task. **(C)** Amount of daily calibration time required for the cursor decoder. **(D)** Time to gain control of the cursor BCI for the first time each day. T15 maintained the ability to calibrate in a few minutes or less over many months. **(E)** Performance of the cursor BCI. T15 maintained the ability to select targets quickly over many months.

T15 configured and toggled cursor control through the BCI’s gaze-based interface, which allowed him to select on-screen buttons to start or stop control, paste recent decoded sentences, and issue function key presses (e.g., escape key; Fig. S6). Each day, he calibrated the cursor and click decoders via a short calibration game. For calibration we used a center-out-and-back target acquisition task with fixed target distance and size until day 621, after which we used a more difficult task with varied target positions and smaller target sizes (see Methods).

Initially, cursor velocity was decoded using a linear model, which required an average of 2.0 ± 1.2 minutes of calibration to gain closed-loop cursor control on a new day (Fig. 3D), and an average of 16.9 ± 7.9 minutes of total calibration per day (Fig. 3C). On day 654, we replaced the linear model with a recurrent neural network (RNN)-based decoder (Fig. S3), which significantly reduced calibration time to just 0.8 ± 0.6 minutes to gain cursor control (rank sum test; p<0.001) and 9.2 ± 5.9 total minutes per day (p<0.001), while maintaining or improving performance (Fig. 3E). This upgrade substantially improved usability, enabling longer, uninterrupted periods of independent computer control.

### Neural signal stability

Speech and cursor decoding performance remained accurate for more than 19 months post-implant, suggesting that the underlying neural signals retained sufficient quality to support high-accuracy decoding. To directly examine the stability of these signals, we analyzed neural activity across the 678-day post-implant period.

Multi-unit action potentials were consistently detected on nearly all electrodes throughout the study. Over 90% of electrodes on each 64-electrode array could reliably detect spiking activity at 2 Hz or higher throughout more than 19-months after implant (Fig. 4A). The dorsal 6v array occasionally exhibited fewer active spiking channels than the other arrays during speaking epochs, consistent with its lower tuning for speech^5^ and stronger tuning for cursor control^18^. Average firing rates for each electrode during speech epochs across the entire dataset are shown in Figure 4B.

**Figure 4.**
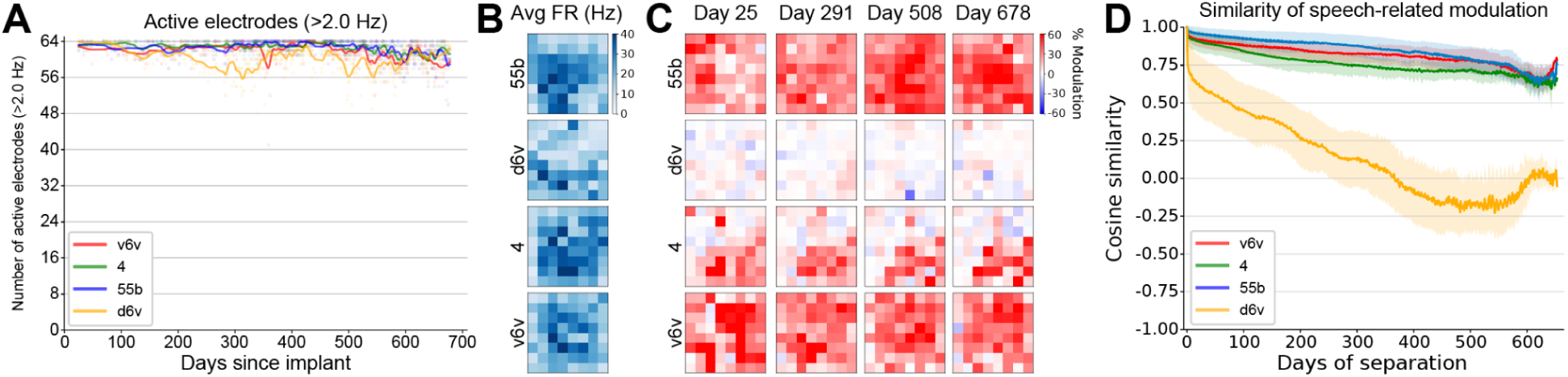
Neural stability. **(A)** Number of electrodes per array with firing rates of 2 Hz or greater during speaking epochs, plotted by post-implant day. Dots are individual data points and lines are Gaussian-smoothed approximations. Colors correspond to the four arrays. **(B)** Average firing rate for each electrode during speaking epochs from the entire 444-session dataset. Electrode locations are drawn according to their physical location on each of the 4 arrays, where up corresponds to dorsal and left corresponds to rostral. **(C)** Speech-related neural modulation for each electrode on a given example day, expressed as the percentage change in spikeband power between rest epochs and speaking epochs. Each column corresponds to the post-implant day indicated by the column title. **(D)** Cosine similarity between daily speech-related neural modulation vectors, as a function of the number of days between each pair. Colors correspond to the four arrays. Solid lines are means and shaded regions represent one standard deviation around the mean.

To assess the stability of task-relevant neural modulation, we measured the change in spike band power between rest and speech epochs for each electrode on each day. These speech modulation maps appeared qualitatively stable across hundreds of days (Fig. 4C). We quantified this by computing the cosine similarity between daily neural modulation vectors for each array as a function of time separation (Fig. 4D). Cosine similarities for all arrays – except the d6v array, which did not exhibit much speech-related modulation – remained above 0.6 even for comparisons separated by more than 18 months, demonstrating that task-relevant neural representations remained quite stable for at least 19 months.

## Discussion

This study demonstrates that an intracortical brain-computer interface can support long-term, independent communication and computer control by a person with severe paralysis and dysarthria due to ALS. Over a 19-month period, the participant used the system in his home for more than 3,800 hours, generating over 183,000 sentences at conversational speeds, browsing the internet, managing personal and professional communications, and maintaining full-time employment. These outcomes establish that a single, speech motor cortex BCI can provide practical, multimodal assistive technology for daily life – enabling both speech and cursor control with high performance and without researcher supervision.

This work successfully addresses two critical gaps that have limited the clinical viability of high-performance intracortical BCIs: the need for researcher-free, at-home operation and the need for sustained decoding performance over long durations. While previous studies have demonstrated high speech or cursor control accuracy in controlled settings, few have supported independent daily use in a home environment, and none to our knowledge have done so for speech or for both speech and computer control simultaneously. Here, the participant and his care partners were able to operate the system independently after an initial training period, and he continued to use it nearly every day, across hundreds of days, over nearly two years.

The system’s stability and usability were made possible by several advances. First, we developed improved architectures for both speech and cursor decoding that reduced calibration time and increased robustness. The speech decoder, built on a transformer-based model, achieved state-of-the-art performance (99.2% word accuracy in Copy Tasks) while requiring little or no daily explicit recalibration from the user. The cursor decoder, upgraded from a linear model to an RNN, similarly reduced calibration time while preserving or improving performance. Both modalities were decoded from the same neural activity recorded in speech motor cortex, where we found sufficient tuning for both speech-articulatory and hand motor imagery. Second, we implemented software features – such as background decoder calibration, gaze-based toggles, and on-screen rating and correction tools – that allowed the system to adapt to the participant’s needs over time, including his shift from vocalized to silent speech. Finally, we automated the system startup and shutdown so that trained care partners could don and doff the system with a few simple steps. We also optimized the system so that, once running, it could operate continuously for up to 19 hours without any care partner intervention.

The participant used this multimodal BCI system as his preferred mode of communication and digital access, instead of existing options such as a gyroscopic head mouse or expertly-interpreted translation through a care partner. He used this system in a variety of personal and professional contexts; the richness and duration of use in this study provide strong evidence that intracortical BCIs can offer not just high peak performance in research settings, but also stable, versatile support for communication and digital interaction during independent home use.

This study has several limitations. It involved a single participant, and the generalizability of these results to other individuals, electrode implant sites, intracortical electrode types, or neurological conditions is not yet known. The system relied on percutaneous wired connections and required daily setup by trained care partners, which may limit broader adoption. We did not systematically assess user fatigue or long-term device wear, although the system’s continued high decoding performance suggests durability across the timescales studied. Future work will be needed to evaluate wireless or fully implantable systems, minimize setup time, and expand access to users with different clinical profiles.

Ultimately, this study demonstrates that stable, high-performance intracortical BCIs can be used independently and productively in a real-world setting, reliably supporting both communication and digital access over nearly two years of use. These findings mark a significant step toward practical, long-term assistive neurotechnologies for individuals with severe motor and speech impairments.

## Methods

### Clinical trial and participant

This study includes data from a single participant (‘T15’) enrolled in the BrainGate2 clinical trial (NCT00912041). This manuscript does not report primary clinical-trial outcomes; instead, it describes scientific discoveries that were made using the data collected in the context of the ongoing clinical trial. BrainGate2 is a sponsor-investigator-led multi-center, open-label, FDA-approved investigational device exemption study (IDE #G090003) assessing the safety of chronically implanted intracortical electrodes (‘Utah’ arrays) in individuals with paralysis. The trial was approved by the Institutional Review Boards at Mass General Brigham (protocol #2009P000505) and the University of California, Davis (protocol #1843264). T15 provided written informed consent to participate in the trial and to publish photographs and video recordings. All procedures were conducted in accordance with relevant guidelines and regulations.

T15 was a man with ALS who was 45 years old at the time of his enrollment in 2023. He retained eye and limited neck movements, but was severely dysarthric. His existing means of communication included communicating through expert interpreters (6.8 WPM) or typing words out letter-by-letter using a gyroscopic head mouse (Quha Zono 2, 6.3 WPM). Further details of T15’s clinical status were previously reported^5^.

In summer 2023, four 64-electrode silicon microelectrode arrays (‘Utah’ arrays; 1.5 mm tine length, iridium-oxide coated; Blackrock Neurotech) were placed in T15’s left precentral gyrus targeting speech motor cortex. Specific information regarding how implant locations were chosen was previously described^5^. Arrays were connected via subcutaneous wires to a percutaneous titanium pedestal affixed to the skull.

### Neural recording and processing

Neural activity was recorded from 256 electrodes using Neuroplex E headstages (Blackrock Neurotech), connected to the two percutaneous pedestals (each linked to two implanted microelectrode arrays). Signals were analog-filtered between 0.3 Hz and 7.5 kHz (4th-order Butterworth), digitized at 30 kHz with 250 nV resolution, and streamed in 1 ms windows to a custom real-time processing node written in Python version 3.8. Each 1 ms window was band-pass filtered between 250 and 5000 Hz using a 4th-order zero-phase Butterworth filter. To reduce edge artifacts, windows were padded using the previous 1 ms of data on the left and mean-padding on the right. Linear regression referencing (LRR)^23^ was applied independently to each array (64 electrodes per group) to suppress common-mode noise.

Two standard neural features were extracted from each electrode: threshold crossing spikes and spike-band power. Threshold crossings were identified when the voltage crossed −4.5 × the root mean square (RMS) value for that electrode. Spike-band power was computed by squaring the filtered signal and averaging over the 1 ms window, with a cap of 12,500 μV² to reject outliers. This full preprocessing pipeline – including filtering, denoising, and feature extraction – was completed in under 1 ms per window. Extracted features were binned into 10-20 ms non-overlapping intervals, depending on the input requirements of each decoder. Binned threshold crossings were computed by summing across consecutive windows; binned spike-band power was calculated by averaging over the same span.

At the beginning of each session, a brief calibration task involving repeated word attempts was used to estimate per-electrode RMS values for thresholding and to compute LRR filter coefficients. These parameters were re-estimated after each block of recording throughout the session to mitigate signal nonstationarities over time.

### Data collection rig

The BCI system was implemented using a distributed, multi-computer setup designed to support both high-bandwidth neural data collection and low-latency, real-time multimodal neural decoding and user interfaces^22^. While the current configuration includes more computational power and physical footprint than would be needed in a future system suitable for widespread clinical use, it enabled rapid prototyping, flexible experimentation, and continuous decoder optimization. In future iterations, the system could be miniaturized into an embedded platform with a portable external interface, similar to prior efforts in reach-and-grasp BCIs using Utah arrays^14^.

Real-time data acquisition and decoding were managed by four networked computers. A Windows 10 machine interfaced with the Neuroplex-E headstages to control neural recording. A second computer (running Ubuntu 22.04 LTS) processed raw 30 kHz neural data, extracted features, and streamed them to a third machine (Ubuntu 22.04 LTS) responsible for GPU-intensive operations including real-time decoder inference, online decoder finetuning, and user-facing task display. A fourth computer (Ubuntu 22.04 LTS) ran the phoneme-to-word language models that generated final text output. All machines were connected via a local area network and synchronized using the Backend for Realtime Asynchronous Neural Decoding (BRAND) framework^22^. Code was implemented in Python, C, and MATLAB.

### Eye tracking

T15’s eye gaze was tracked using a Tobii Pro Spark eye tracker (Tobii AB, Stockholm, Sweden) that was mounted at the bottom of the BCI system participant monitor. Calibration was performed at the start of each session (for both research sessions and independent sessions) and repeated as needed via an option available in the BCI system menu. Gaze location was sampled at 60 Hz, averaged across both eyes, smoothed over time, and used to enable gaze-based selection of on-screen buttons. Button activation was triggered by maintaining gaze on an on-screen target for 0.5-1.0 seconds.

To improve usability and reduce selection errors, we implemented a custom “magnetization” feature that subtly attracted the gaze cursor to the center of nearby on-screen buttons, making them easier to select. Eye tracker calibration, gaze data acquisition, and selection logic were implemented in custom Python software integrated into the BRAND-based data collection platform.

### Data collection sessions

All data for this study was collected in one of two types of data collection sessions: scheduled research sessions or personal use sessions. All data collection sessions occurred in the participant’s home. Scheduled research sessions occurred 0-2 times per week, and during these sessions the participant would be asked to do a research-focused task that, for the purposes of this study, most often involved attempting to say prompted sentences aloud in a structured Copy Task. By contrast, personal use sessions had no research-focused tasks other than what was necessary for enabling use of the BCI for the participant’s self-directed communication and digital access. Prior to post-implant day 281, personal use sessions were scheduled 0-4 times per week and required a member of the research team (co-author C.I.) to be present to don and doff the system. After post-implant day 281, following an amendment to the Investigational Device Exemption, T15 and his care partners could use the BCI system whenever they wanted to without researcher assistance. Data from all sessions, except for when “Privacy Mode” was enabled during personal use sessions, was saved and used to train future speech decoding models.

### Speech decoding

#### Neural feature preprocessing

Neural activity was decoded into text using two types of neural network architectures used over the course of the study: a recurrent neural network (RNN) and a transformer-based model. Both architectures processed 512-dimensional neural feature vectors (threshold crossings and spike-band power from 256 electrodes, binned every 20 ms), which were z-scored using statistics from the preceding 5-20 speech trials and smoothed using a Gaussian kernel (σ = 40 ms).

#### Phoneme classification

Both models were trained to predict a distribution over 41 output classes: the 39 standard American English phonemes plus a “silence” class and a CTC blank token. This formulation enabled alignment-free training using the connectionist temporal classification (CTC) loss function^7^. Sentence labels for each trial were converted to phonemes using the CMU pronunciation dictionary^24^ and the *g2p-en* Python package^25^.

#### RNN decoder architecture and training

A recurrent neural network (RNN) predicted phoneme probabilities from neural activity. The model consisted of three components: (1) a day-specific linear input layer to account for inter-session neural drift; (2) five stacked GRU layers, each with 768 units; and (3) a dense output layer producing a distribution over 41 classes. The RNN produced outputs every 80 ms based on the most recent 280 ms (14 bins) of neural data.

Offline training was performed prior to each session using all previously collected data. Training used 90% of past trials and randomly held out 10% per day for validation. Each batch (up to 64 trials) was sampled from one session, with real-time data augmentation including white noise and constant offsets. Optimization was done with Adam (learning rate linearly decayed from 0.02, β_1_=0.9, β_2_=0.999, ε=0.1), with dropout and L2 regularization.

#### Transformer decoder architecture and training

The transformer-based decoder consisted of three stages: day-specific transformation, input striding, and a 12-block causal transformer with multi-head attention. Day-specific embedding input layers added a learned day-specific vector to each input timestep to correct for inter-day variability. This linear offset (as opposed to a nonlinear transformation as was used for the RNN^5,7^) was empirically found to perform better for this architecture. The transformer decoder applied multi-headed causal attention with Rotary Positional Embeddings^26^ and used RMSNorm^27^ for normalizing hidden states. Outputs from the transformer were fed through a feedforward layer to produce phoneme probability distributions every 80 ms. Like the RNN, this model was also trained with CTC loss, which automatically aligned neural data and labeled phoneme sequences.

Offline model training used all previously recorded labelled data (i.e. Copy Task trials and personal use trials that were confirmed by the participant to be decoded correctly), excluding trials that were too noisy or too long, with a 90/10 train/validation split per day. Data were augmented with noise and constant offsets, and then smoothed using a Gaussian kernel (σ = 40 ms). Mixed-day batches (6 random days of data, 16 randomly-selected trials each) were used to regularize learning. Optimization was performed with AdamW using a cosine decay schedule and warmup. Training lasted for 400,000-600,000 batches and was run on an RTX 4090 GPU.

#### Continuous finetuning (both models)

During real-time use, both RNN and transformer models underwent continuous finetuning. Once a minimum number of new sentences were collected (5-6), a new day-specific embedding was created. Each new sentence had a 67% chance of entering the training buffer and 33% chance of being held out for validation. Finetuning batches (size 64) drew data from the current and three random prior days. Checkpoints were updated only if validation performance improved. Finetuning occurred in 10-batch intervals for up to 500 batches per session using cosine learning rate decay. This continual adaptation improved decoding stability and accuracy by tracking nonstationarities across sessions.

### Cursor decoding

#### Calibration task

Until post-implant day 621, the calibration task was a center-out-and-back eight target “Radial 8” acquisition task with fixed target distance (40% of screen height), fixed target radius (5% of screen height), and the next target predictably in the center after every outward trial. After day 621, the task was made more varied and engaging to elicit more diverse cursor movements (similar to the previously reported Fitts task ^28^). Target positions could be anywhere in the 2-D workspace (1920 x 1080 pixels), leading to both shorter and longer target distances, and cursor trajectories at any angle. Target radii were smaller than in the Radial 8 task and varied (2.5-4.0% of screen height).

#### Decoder architecture and training

Cursor control originally used a linear velocity decoder as reported previously^18^, which T15 calibrated anew each day. After post-implant day 467, the decoder’s speed gain and nonlinear speed adjustment cutoff were set to new values which improved T15’s ability to stop on and click small buttons during computer use. This improvement was reflected in a high-performance grid evaluation task which inherently rewards precision^9,18,29–32^, but was not reflected in the performance on the calibration task (Fig. 3E), whose larger targets tolerated faster cursor movements at the cost of precision.

After day 654, cursor control used an RNN-based decoder. The RNN was trained offline on up to 44 prior sessions of cursor calibration data. Unlike the linear decoder, the RNN was not updated online during calibration. However, the calibration task was still used daily to train a click decoder and to collect up-to-date input feature statistics (spike band power means and standard deviations) for input normalization.

Each input to the RNN was a 256-D vector of z-scored spike band power for a 10 ms time bin. Every 10 ms, the decoder output a 2-D vector {𝑥, 𝑦} representing a predicted cursor velocity, and then moved T15’s cursor accordingly. The RNN model consisted of a two-layer GRU with a hidden size of 64, followed by a dense layer projecting the hidden state to a 2-D output vector (Fig. S3).

During training, labels were 2-D vectors from cursor to target (“target vector”). The target vector was set to {0, 0} when the cursor was touching the target under the assumption that the participant was trying to hold still^33,34^. Training aimed to minimize the per-sample loss function

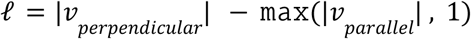

where 𝑣_𝑝𝑒𝑟𝑝𝑒𝑛𝑑𝑖𝑐𝑢𝑙𝑎𝑟_ is the component of the predicted velocity 𝑣 that was perpendicular to the target vector and 𝑣_𝑝𝑎𝑟𝑎𝑙𝑙𝑒𝑙_ is the component of the predicted velocity 𝑣 that was parallel to the target vector. This loss function encouraged output vectors that pointed toward the target, with a bounded magnitude. This loss function also encouraged output vectors of {0, 0} while the cursor was touching the target. Each training epoch, Gaussian noise was added to both the inputs (z-scored spike band power) and labels (target vectors), as a form of data augmentation^35^.

During online cursor control, output vectors were scaled by a speed gain β, which T15 could raise or lower using the on-screen user interface.

### Click decoding

Click decoding used a linear decoder as reported previously^18^, which T15 calibrated anew each day as part of the same tasks used to calibrate cursor control. The decoder’s weights were computed online using logistic regression every 3.0 seconds during calibration.

Each input to the click decoder was a 512-D vector of z-scored neural features (256 threshold crossings values and 256 spike band power values) for a 10 ms time bin. Every 10 ms, the decoder output a discrete class “click” or “no click”, and the BCI performed a left-click on T15’s computer for each “click” decoded.

To reduce errant clicking, a “click” was only decoded when the linear model’s probability estimate for “click” was higher than for “no click” for some minimum portion 𝑇 of time bins within a 70 ms window. T15 could raise or lower the click sensitivity (1 − 𝑇) using the on-screen interface.

### Decoder evaluations

#### Speech decoding

Consistent with prior studies^5,7,8,10,11^, we benchmarked our speech decoding accuracy by calculating the predicted phoneme and word error rates during a structured Copy Task where the participant was asked to say prompted sentences aloud, thereby allowing us to directly compare the decoded output with the ground truth target sentence. Word and phoneme error rates were calculated using Levenshtein distance, which counts the number insertions, deletions, or substitutions necessary to match the decoded phonemes or words to the ground truth labels. Reported error rates were aggregated across all evaluation sentences from each session by summing the number of errors (insertions, deletions, or substitutions) for all sentences and then dividing it by the total number of words in those sentences. This helps prevent very short sentences from overly influencing the result. Confidence intervals for error rates were computed via bootstrap resampling over individual trials and then re-calculating the aggregate error rates over the resampled distribution (10,000 resamples).

During independent BCI use, we did not know the ground truth of what T15 was saying, and hence we could not compute the true phoneme and word error rates. Instead, we asked the participant to use gaze or cursor control to label the accuracy of each decoded utterance as “100% correct”, “one word wrong”, “mostly correct”, or “incorrect”. He also had the opportunity to make corrections to decoded sentences before rating the accuracy. We instructed T15 on the importance of proper labeling of the sentences and encouraged him not to under- or over-report the decoded outcome. We shared with him that the proportion of “correct” sentences would be further used for the recalibration that was necessary for ongoing system maintenance, and that incorrect sentences would be used to help troubleshoot our system for future software iterations. Hence, he was incentivized to correctly label outcomes to maintain accurate ongoing use. When asked how he chose between labeling a sentence with multiple incorrectly decoded words as “mostly correct” or “incorrect”, T15 told us that he uses a threshold of 2/3rds (Table S1): sentences with more than 2/3rds of words decoded correctly are “mostly correct”, and sentences with less than 2/3rds of words decoded correctly are “incorrect”.

#### Cursor decoding

Formal evaluations of cursor decoding performance were done in structured research sessions via a Grid Task^9,18,29–32^, from which we could compute bits per second. During daily independent BCI use, however, the only structured cursor-related tasks that the participant performed were the center-out-and-back or random target acquisition tasks, which were used to calibrate the cursor decoder. Thus, we relied on metrics from these tasks to quantify the daily cursor decoding performance. These measures included (1) how long it took the participant to achieve full closed-loop cursor control during the calibration task, (2) the target acquisition rate during the calibration task, and (3) the total minutes of calibration per day.

### BCI system software (user interface)

We built a custom user interface that supported the daily independent use of the BCI for speech decoding, cursor control, personal computer control, and a variety of related options. The user interface was primarily coded with the *Pyglet* Python package version 2.0.12^36^ and consisted of a range of function-specific “pages” that the user could navigate between using gaze or cursor control (Fig. S6). This interface was displayed on a monitor that was mounted on an articulating arm, which was positioned above the participant’s personal computer (e.g., see Fig. 3A). The system was periodically updated in response to participant feedback, feature requests, and bug fixes in the underlying custom software stack (Table S2).

While idle, the system waited to either detect attempted speech or a user interface function press via gaze or cursor. During attempted speech, predicted words were displayed on screen in real time. After the participant was done speaking, he could optionally use the interface to make corrections to the decoded word sequence before rating its accuracy. At any time, the participant could have the most recently decoded sentence played as audio via text-to-speech or typed into the active text field on his personal computer. He could also navigate to a menu screen, which included a variety of options including cursor calibration, eye tracker calibration, the ability to toggle personal computer cursor control on or off, cursor-related options such as move speed or click sensitivity, a toggleable profanity filter, a toggleable privacy mode wherein data would not be logged or saved in any way, a full gaze or cursor-controlled keyboard, and more. Snapshots of the full BCI system user interface are shown in Fig. S6.

### BG Home software (personal computer integration)

A custom application (‘BG Home’) enabled our BCI system to control the participant’s computer’s mouse and key presses such that he could use his personal computer via BCI control. It consisted of a small user feedback window hovering in the corner of the computer screen, plus an asynchronous background process which received streaming data from the BCI system and performed the mouse movement and key presses. BG Home was built with Electron, but used Python for the background process. A snapshot of what the BG Home software looks like while the participant is attempting to speak can be seen in Fig. 3A.

## Extended figures

**Figure S1.**
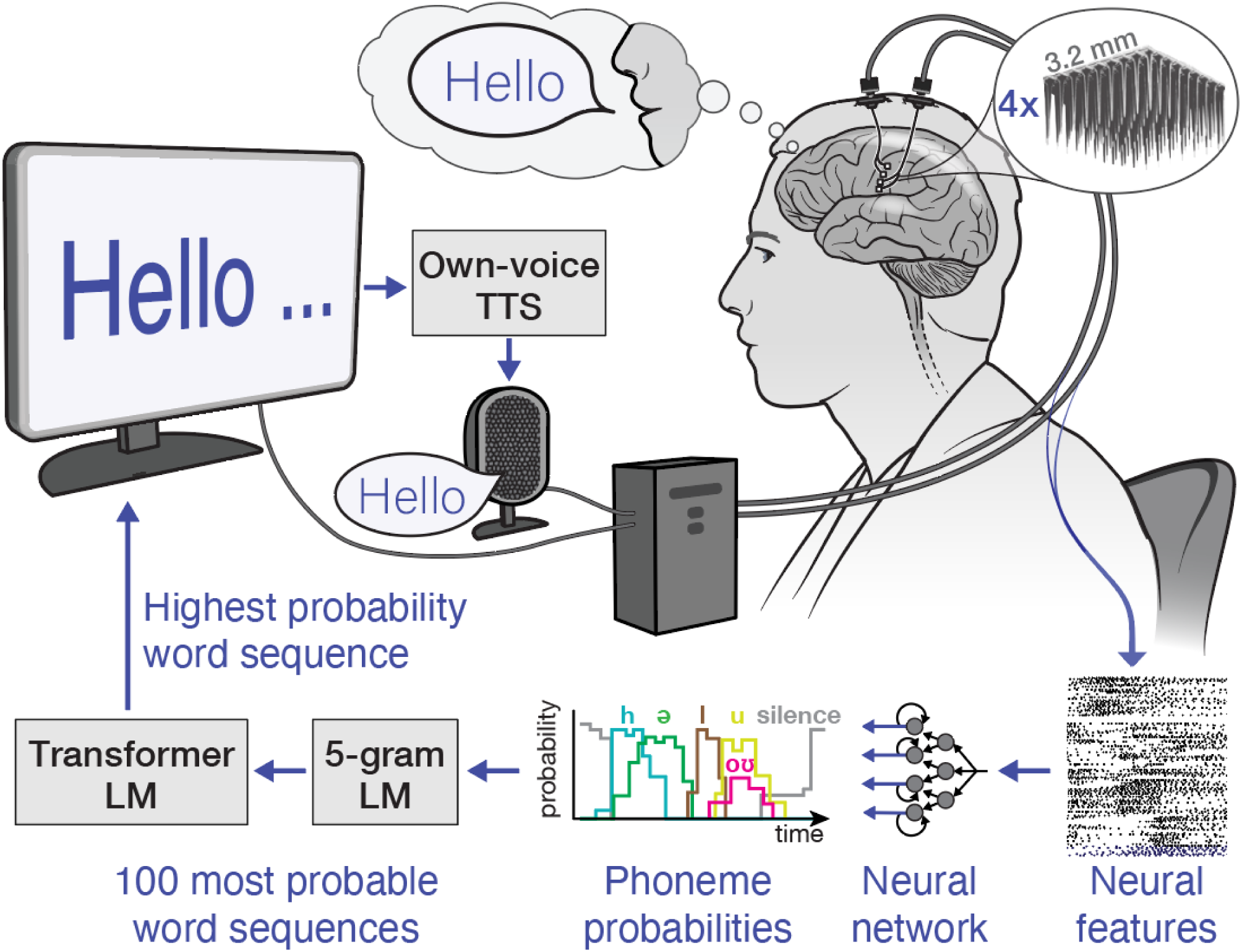
Brain-to-text decoding pipeline. Neural signals are recorded from the speech motor cortex via four 64-electrode ‘Utah’ arrays as the participant attempts to speak. The neural activity is processed in real time into neural features, which are inputted into a neural network to predict the most likely phonemes that the participant is trying to say every 80 ms. Phoneme probability sequences are inputted into a multi-stage language model to identify the most likely word sequence being spoken. Predicted words are then displayed as text on a display in real time, and at the end of a sentence they are read aloud by a text to speech (TS) model that was optimized to sound like the user’s pre-ALS voice.

**Figure S2.**
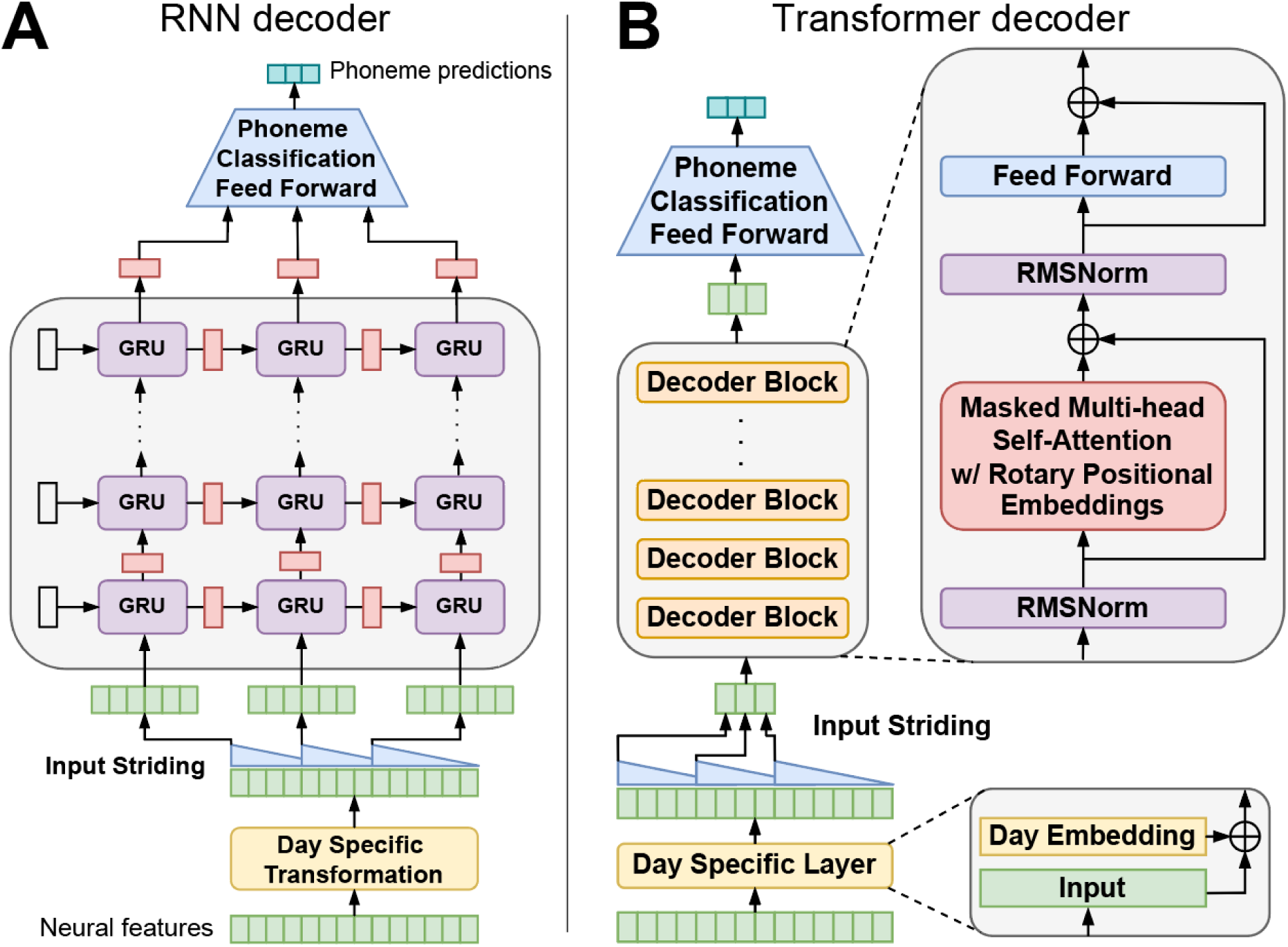
Brain-to-text decoder architectures. **(A)** A recurrent neural network (RNN) based neural network that predicts the most likely phoneme being spoken from sequences of neural features. Neural features are inputted to the model and then undergo a day-specific transformation with the goal of aligning nonstationary neural data from separate days into a common subspace. Aligned features are then rearranged into overlapping time windows (“input striding”) before being passed to a series of GRU layers. Finally, a feedforward layer converts the output to the probability that any English phoneme is being spoken. This model was trained with connectionist temporal classification (CTC) loss, which automatically handles alignment between neural data and phoneme labels. **(B)** A transformer-based neural network that predicts the most likely phoneme being spoken from sequences of neural features. The inputs, outputs, and loss function of this model are identical to the RNN model in (A), but the transformer-based internal architecture is different. We implemented multi-head self-attention and applied causal masking so that the model can be used in a streaming application. We used Rotary Positional Embeddings for positional encodings and RMSNorm for normalizing activations before attention layers and feedforward layers.

**Figure S3.**
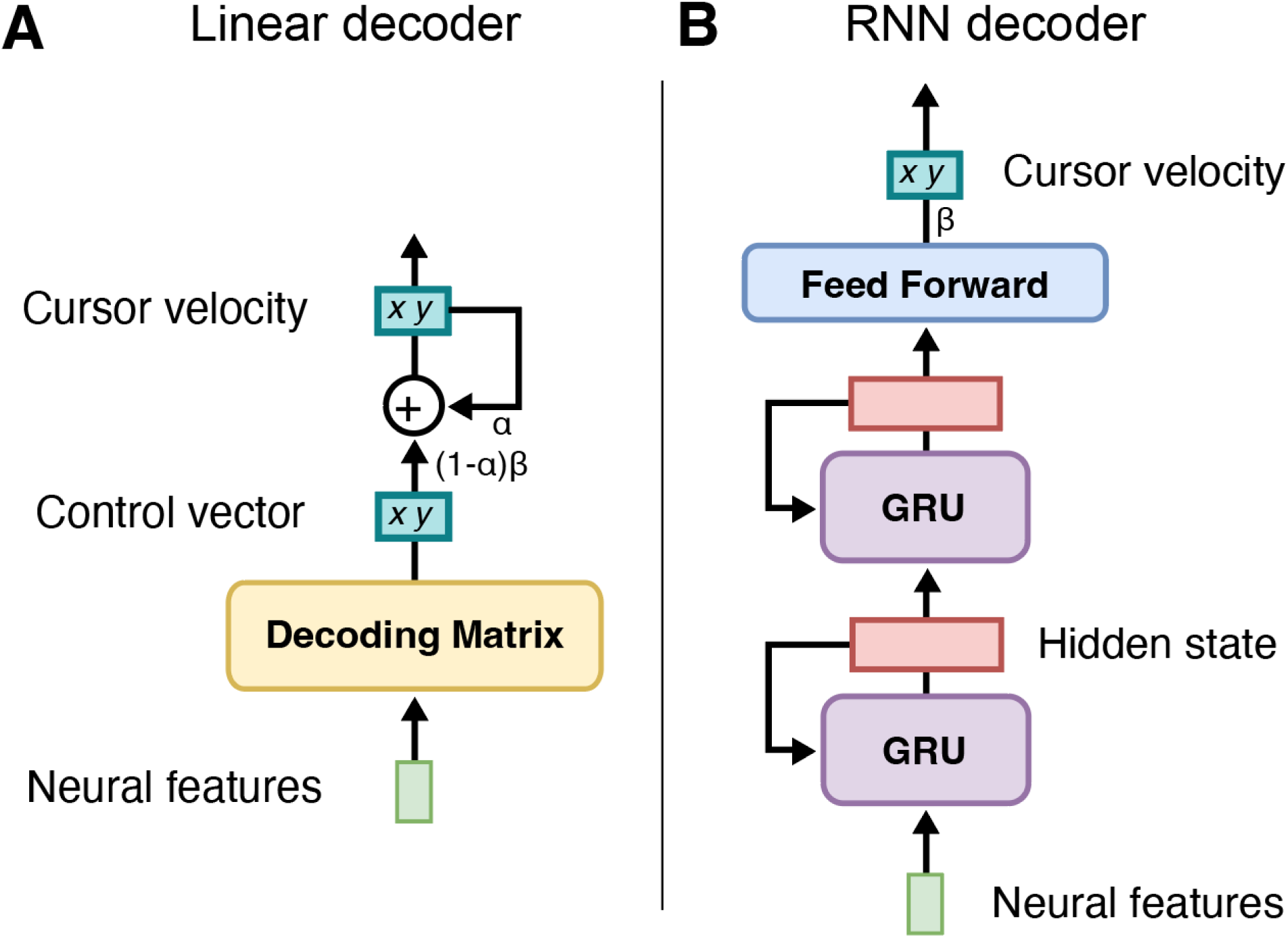
Cursor decoder architectures. **(A)** A linear velocity decoder with smoothing, for decoding intended cursor velocity from neural features. For each 10 time bin, a decoding matrix projects a vector of neural features into a 2-D control vector (the ‘dimensionality reduction’ step). This is averaged with the previous time bin’s predicted velocity (the ‘smoothing’ step) where α sets the balance between the current control vector and the previous velocity and β modulates the overall speed. **(B)** A recurrent neural network (RNN) based neural network for decoding intended cursor velocity from neural features. For each 10 ms time bin, a vector of neural features is inputted to a 2-layer GRU model. A feedforward layer transforms the output of the GRU into a 2-D velocity vector. β modulates the overall speed. The hidden state from each GRU layer is passed back into the model for the following time bin.

**Figure S4.**
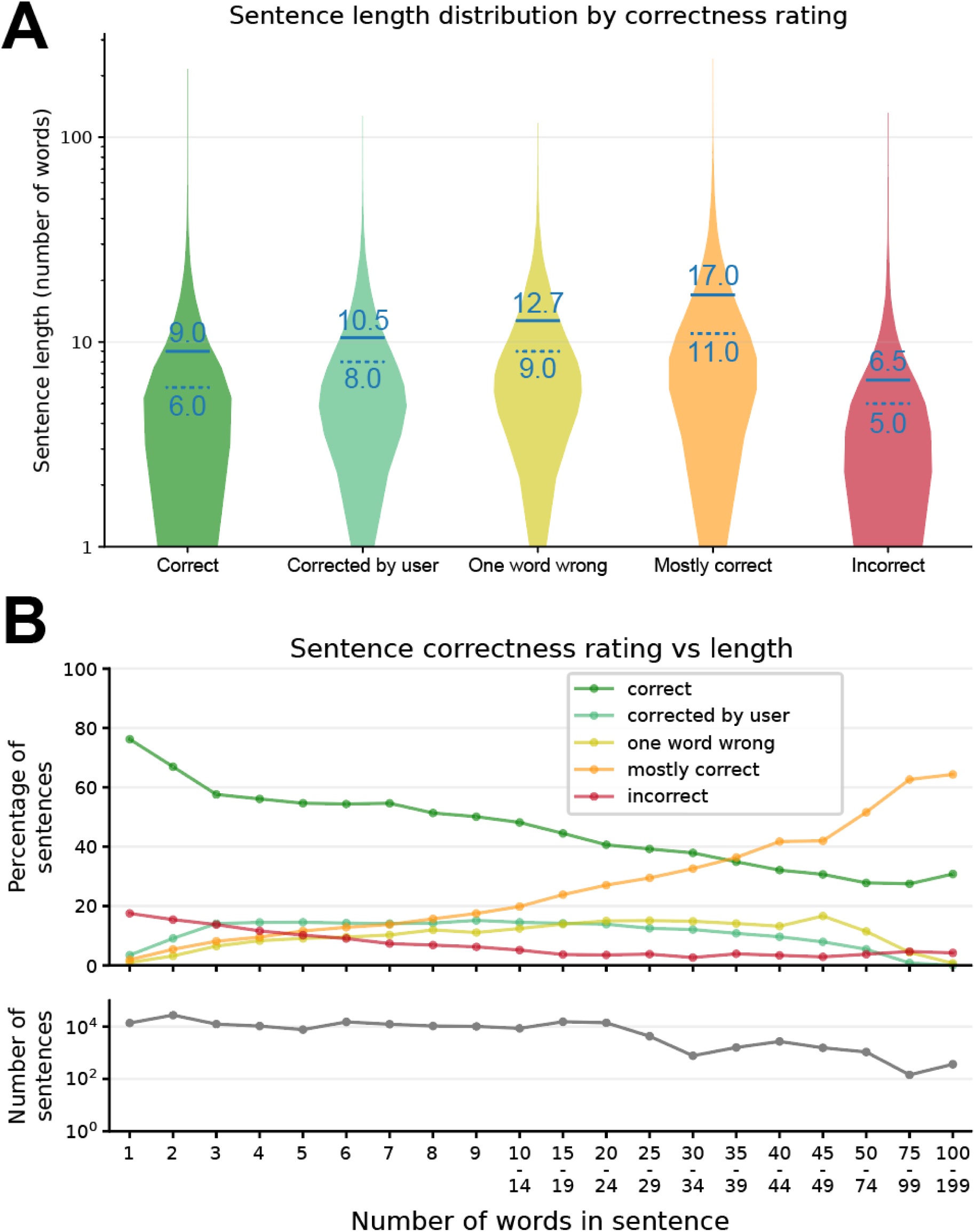
Sentence length versus decoding accuracy. **(A)** Cumulative distributions of sentence lengths as a function of participant-rated decoding accuracy. Solid horizontal blue lines are means, dashed lines are medians. **(B)** Sentence correctness rating probability displayed as a function of sentence length.

**Figure S5.**
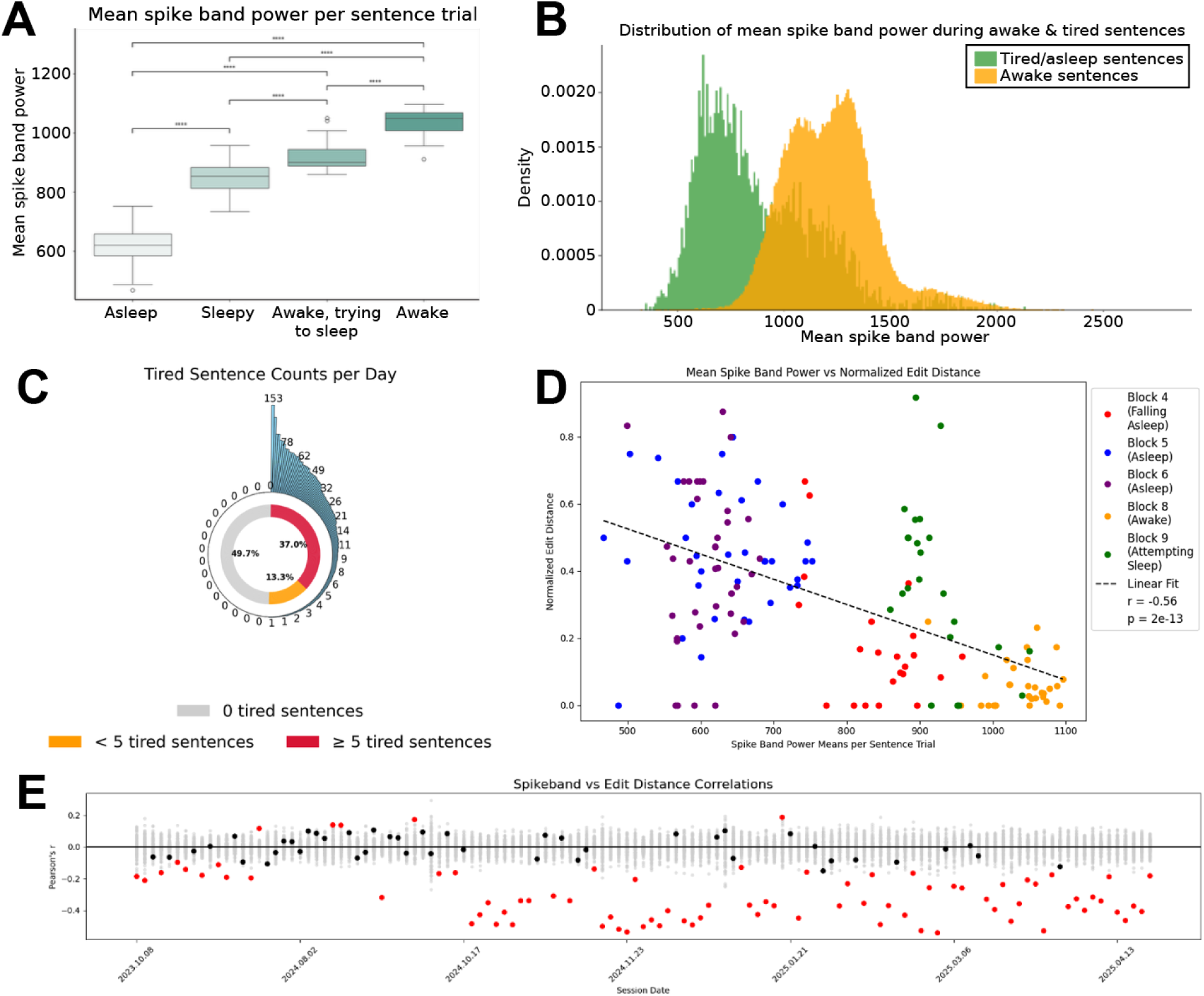
Participant’s level of alertness affects neural signals and speech decoding accuracy. **(A)** Relationship between mean spike band power and alertness level. Neural data were recorded in a structured research session while the participant was asleep, sleepy, awake but trying to sleep, and awake. Spike band power varied significantly with alertness level (ANOVA with post hoc comparisons; ***p<0.001). **(B)** Average spike band power during independent BCI use while the participant was alert versus tired/asleep. Each datum represents the average spike band power for a sentence; data from all personal use sessions where eye tracking data was available are aggregated, resulting in 139,364 alert trials and 3,714 tired/asleep trials. Alertness level was inferred using eye tracking data: if no eye tracking data was available for 2 minutes or more, alertness was categorized as tired/sleeping. **(C)** Number of daily sentence trials where the participant was tired or sleeping per day of independent use. 37.0% of days had at least 5 sentences that were categorized as tired. **(D)** Relationship between average spike band power and speech decoding accuracy. Here, speech decoding accuracy is quantified as the edit distance between the predicted phoneme sequence output from the RNN/Transformer neural decoder and the predicted phoneme sequence output from the language model – higher values indicate worse speech decoding. Points are individual trials, colored according to the participant’s alertness level. A linear trendline was fit to the data, which had a significantly negative slope, indicating that speech decoding tended to be better on trials with larger average spike band power values where the participant was more alert. **(E)** Linear correlation values between mean spike band powers and edit distance per trial across each day of independent use. Chance correlations were estimated with 100 shuffled datasets’ values and shown in light gray. Significant correlations (p < 0.01) are marked in red.

**Figure S6.**
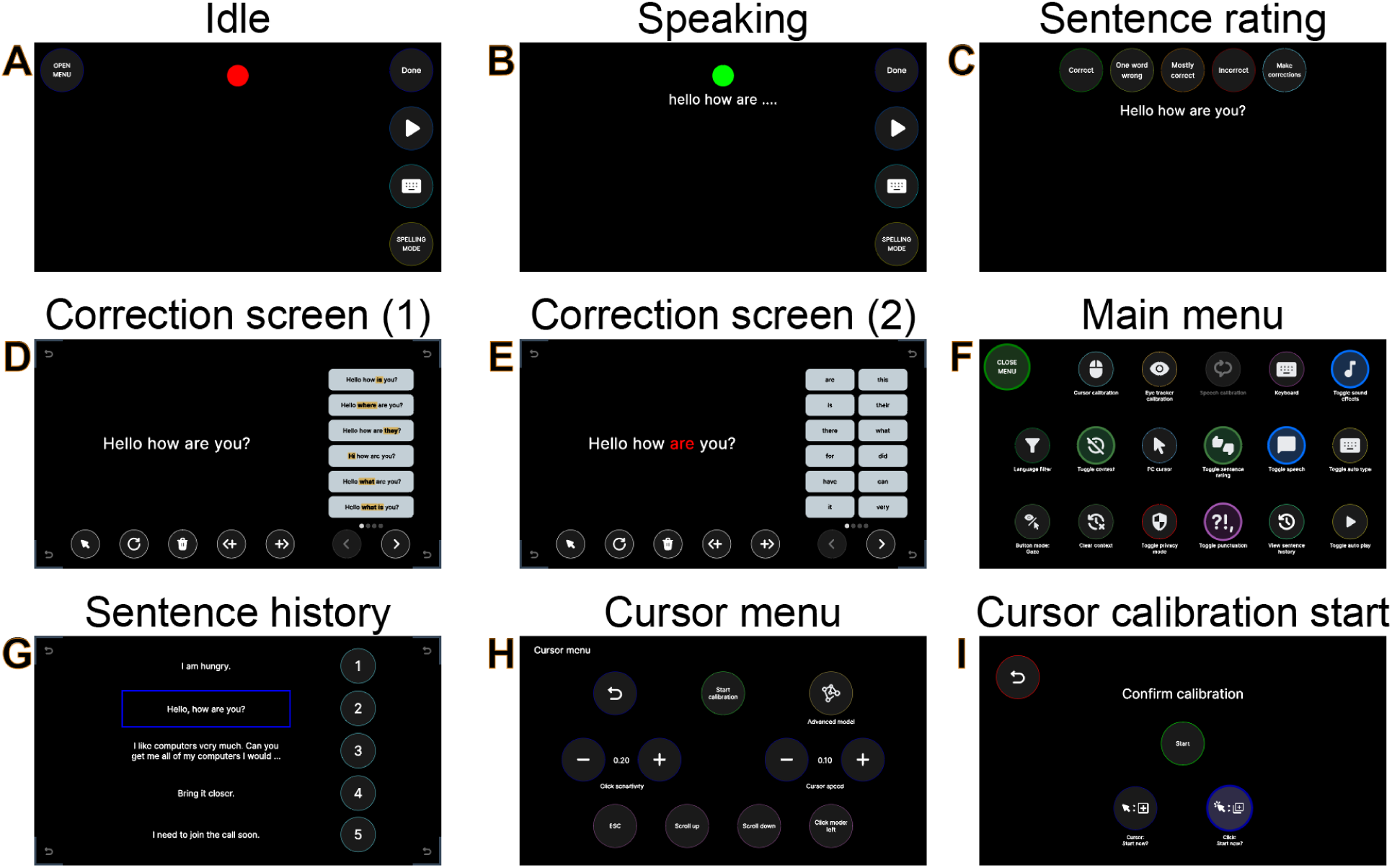
BCI system user interface. **(A)** Graphical user interface (GUI) during idle epochs, where the participant is not speaking. On-screen buttons can be selected with gaze or cursor control. The “OPEN MENU” button opens the menu screen (F). The “DONE” button manually stops speech decoding when pressed. The button with a play symbol (▶) plays the most recently decoded sentence aloud using text-to-speech, and the keyboard button pastes that sentence into the active text field on the participant’s personal computer. The “SPELLING MODE” button triggers spelling mode, where the language model vocabulary is restricted to the 26 English letters rather than the normal ∼125,000-word vocabulary. **(B)** GUI during speaking epochs. When speech is detected, the center circle turns green, decoded words appear on-screen in real time, and the menu button is hidden. When the user is done speaking, they can press the “DONE” button to manually end the sentence or they can wait for a 6-second timeout, after which the sentence will be automatically ended. **(C)** GUI during sentence rating, which occurs at the end of every sentence unless the feature is turned off in the menu by the user. The user can choose to rate the accuracy and move on, or to make corrections to the sentence using the interfaces in (D) and (E). **(D)** GUI for making corrections to incorrectly decoded sentences. Here, no incorrect word is selected so alternative sentences are shown on the right side, with word differences highlighted in yellow. Corner revert-icon buttons allow the user to exit this screen. The circular buttons on the bottom, from left to right, are: toggle between gaze control and cursor control, refresh word suggestions, delete selected word, insert new word to the left of selected word, insert new word to the right of selected word, page forward, page back. **(E)** Same as (D), except now an incorrect word has been selected by the user (red font) and alternative words are shown in the list on the right. **(F)** Main menu screen. Here, the user can enable or disable various functions or enter submenus by selecting each option. The button in the top left exits the menu back to the idle screen as in (A). **(G)** Sentence history screen, entered using the corresponding button in the menu in (F). Recently decoded sentences are shown and can be selected for playing through TTS or pasting into the user’s personal computer. **(H)** Cursor menu screen, entered using the corresponding button in the menu in (F). Here, the user can toggle between the linear or RNN-based cursor decoder, change cursor speed and click sensitivity, toggle presses of function keys for their personal computer, and enter the cursor calibration game. **(I)** Confirmation screen before the cursor calibration game begins. The user can decide whether to update the cursor decoder, click decoder, or both.

**Table S1.** Questions and answers with T15.

| Question | T15's answer |
| --- | --- |
| Please briefly explain your rationale for how and when you rate the accuracy of decoded sentences. The “100% correct” and “one word wrong” ratings are self-explanatory, but how do you decide between rating a sentence “mostly correct” or “incorrect”? For example, a 10-word sentence that has 2 words wrong might be “mostly correct”, but a 3-word sentence that has 2 words wrong might be “incorrect”. | I use a percentage of two thirds to assess whether something is good enough to be mostly correct. It needs to be that or higher. |
| How frequently do you turn off the sentence rating feature, and what are the typical reasons for doing so (general reasons are fine, I don't need specifics)? | Probably about three or four times a week, and always for something connected to physical comfort. I may be moving my chair too far away to have access to the eye tracker technology, so I will do this for simplicity. It will save me time, or alternatively, I will want to be able to talk very quickly for emergencies. I will turn it off for this reason. But likely less than one percent of my overall sentences and such. |
| When a sentence is not decoded completely correctly, how do you decide whether to enter the correction interface or to skip it and mark the sentence as “one word wrong”, “mostly correct”, or “incorrect”? | I would normally check whether a one mistake sentence is able to get changed into a one hundred percent correct sentence. And after you told me how important this is, I have been checking every time. On the other hand, if the sentence is completely hot garbage, I will label it incorrect before I check it. The longer the sentence, the more likely it will be eighty five percent or higher correct. Most sentences that are more than ten words and are labeled mostly correct are in between eighty five to ninety five percent correct. FYI I am usually only skipping the sentences that are obviously missing the right count of phonemes. The decoder will get off track and the sentence will be completely ruined. |
| When you choose to enter the correction screen, roughly what percentage of the time do you think you are able to successfully correct the sentence? | I think I know what you are looking for here. I am always almost able to correct a word or two in a sentence with multiple mistakes. But I think that I am able to get a sentence all the way to one hundred percent correct about thirty to forty percent of the time. |
| Since we implemented the upgraded brain-to-text decoder [on post-implant day | Yes, it is difficult to give quantitative feedback about this, but the answer is a resounding yes. I would say the amount of sentences that are one hundred |
| 602], have you noticed improved or more reliable decoding accuracy? | percent correct has improved, as well as the number of hard to decode words being correctly decoded. |
| Do you notice any trends between decoding accuracy and (a) how you are feeling or (b) what you are trying to talk about? | I think that it is a relatively narrow and very high range of different days I experience in terms of decoding accuracy. So, while the answer is yes, it is not something that I am saving a particular conversation for when the decoder is having a good day. Being able to fully communicate to people is a dream. We are not there yet, but this is such a massive step forward and every day that we can improve something, even a little tiny bit is felt by me as something really positive and will impact my mood and my day, absolutely. |
| How has your overall experience with personal use been in the last few months? How does it compare to last year? | It feels like we are still on the upward swing. With a dedicated team of people working on this, we are still making improvements on the utility of this service. Every day is a new challenge to solve, but I feel like we are only getting better. This would be very different if I was going down hill and losing my ability to talk. |
| Broadly, what do you typically use our system for throughout a day, in terms of controlling your own personal computer? For example, could you estimate the number of text messages (texts, slack, emails) you might send in a day, or whether you use it for video calls, or just to relax and watch a show or search the web? | I will send a lot of very short messages on multiple points during the day. Probably twenty to thirty one line messages. slack. signal. text. email. Then I would probably have two or three longer messages, like this email that I will send in a day. Or I will make a similar contribution on a Google document or sheet. Definitely will get in a video or two as a little treat. I will use the device to tell [my assistant] what to do and give him feedback. During the work week, we will definitely hop on a few calls every day. And on the weekends or the evening, I will have visits from friends and family, or usually I will call with them. I will use the internet to get everything because I do not usually leave my house. So that is incredibly important to be able to stay on top of my finances and shopping. It will take me more time, but I am basically using my computer in the same way as I was doing before the disease. |
| Any other comments you want to share that my questions did not cover? | Look at the very small or short sentences, about three words or less, and you know that the decoder will work much better with more context. And I would think that the majority of completely incorrect sentences are coming from my obsession about |
|  | trying to get the exact right word. If you take those sentences out, you will get very different data. |

**Table S2.** Timeline of updates and changes to the BCI system.

| **Post-implant day** | **Feature** | **Description** |
| --- | --- | --- |
| 147 | Bluetooth keyboard | Enables the BCI system to act as a bluetooth keyboard for the participant's personal computer so that they can type decoded text into text fields on their computer. |
| 227 | Sentence correction | Enables the user to choose between the top 5 most likely decoded sentences if the top sentence was not decoded correctly. |
| 232 | Eye tracker button magnetization | "Magnetizes" on-screen eye tracker buttons so that tracked gaze will gravitate toward them when nearby. This makes gaze-based control easier to use and more robust to imperfect calibration. |
| 281 | Independent mode | System was automated so that care partners could power it on and off without researcher assistance. |
| 295 | BG Home version 1 | Version 1 of the BG Home software suite was deployed, which integrates the BCI system directly with the user's computer. |
| 303 | Added "one word wrong" sentence rating option | At the participant's request, we added the "one word wrong" sentence rating option alongside the existing "100% correct", "mostly correct", and "incorrect" options. |
| 358 | Enabled cursor control for personal computer | This update involved adding a new menu screen to the BCI system UI, a cursor calibration game, and a major update to the BG Home software that supported controlling mouse inputs. |
| 361 | Use recent context to help predict the current decoded sentence. | Recently decoded sentences are fed as context into the language model alongside the current most likely candidate sentences to help choose the most contextually appropriate option. Recent context is also displayed on the screen to help conversation partners keep up with the current topic of conversation. |
| 364 | Eye tracker calibration | Enabled the participant to initiate an eye tracker calibration game via the gaze- or cursor-based user interface without care partner assistance. |
| 412 | Speech decoder background calibration bug fix | A major bug was identified and fixed in the background calibration code for the speech decoder, which resulted in immediately increased speech decoding accuracy during personal use. |
| 462 | Word-based correction | Instead of making corrections by choosing among the top 5 most likely decoded sentences, the participant can now choose individual words that were decoded incorrectly and replace them by choosing from a list of contextually appropriate replacement words. |
| 462 | Privacy mode | The participant can enable a privacy mode via the gaze- or cursor-based interface. When enabled, no data will be saved. |
| 467 | Optimized cursor parameters | The cursor decoder's speed parameters were set to |
|  |  | more optimal values for better precision (selecting tiny buttons) and traversal (moving across the screen). |
| 484 | Miscellaneous updates | Visual user interface refinements, a bad-word language filter, keyword sound effects, ability to disable speech detection, ability to disable sentence rating. |
| 486 | Optional cursor control on BCI system user interface | Instead of relying on the eye tracker exclusively, the participant can now also use cursor control to navigate the BCI system user interface. |
| 491 | Refresh word suggestions | Enabled the participant to generate new updated word suggestions during word correction mode. This is useful for when he makes some corrections, then can refresh to get more contextually relevant options before continuing to make additional corrections. |
| 505 | Word insertion and deletion | Enables the participant to insert or delete words from the decoded sentence during word correction. |
| 561 | Speech detection animation | Added a user interface element that animates (grows and shrinks in size) whenever speech is detected by the speech decoder. |
| 563 | Care partner UI assistance | Added the ability for a care partner to use a physical mouse to navigate the BCI system user interface in the event that the participant needs help (e.g., if gaze control stopped working and needs to be recalibrated). |
| 565 | Gaze- or cursor-based full keyboard | Added a full keyboard that can be controlled with gaze or cursor so that the participant can type things on his personal computer that would be difficult or impossible to do through the speech decoder (e.g., a website address). |
| 565 | Automatically pause computer cursor control during gaze | When the participant's gaze is detected on the BCI system screen, computer cursor control is paused so that errant cursor movements or clicks do not occur. |
| 588 | Aesthetic user interface changes | Font was changed throughout the system. The gaze selection circle on the word correction screen was animated to make it more clear when a dwell selection was complete. |
| 602 | New transformer-based brain-to-text decoder | A transformer-based brain-to-text decoder was deployed, resulting in higher speech decoding accuracy with less calibration required. |
| 612 | Cursor- or click- only calibration modes. | The participant can now choose whether to calibrate the cursor decoder, click decoder, or both. Prior to this change, both decoders always had to be calibrated in tandem. |
| 616 | Updated word correction screen UI | Updated and optimized the user interface for the word correction screen to provide a more seamless user experience. |
| 621 | New cursor calibration game | Switched from a center-out-and-back style cursor calibration game to a varied target task to get more diverse cursor training data. |
| 624 | More word correction options | During word correction, the participant is now offered a wider variety of potential replacement words. |
| 651 | Refactored user interface code | Refactored the entirety of the BCI system user interface code, resulting in higher frame rates and a variety of bug fixes. |
| 654 | Upgraded to RNN-based cursor decoder | Upgraded the cursor decoder from a linear-based model to an RNN-based model, resulting in similar or improved performance with less calibration time. |

## References

1. Coppens P. Aphasia and Related Neurogenic Communication Disorders. Jones & Bartlett Publishers; 2016.

2. Katz RT, Haig AJ, Clark BB, DiPaola RJ. Long-term survival, prognosis, and life-care planning for 29 patients with chronic locked-in syndrome. Arch Phys Med Rehabil 1992;73(5):403–8.

3. Lulé D, Zickler C, Häcker S, et al. Life can be worth living in locked-in syndrome [Internet]. In: Laureys S, Schiff ND, Owen AM, editors. Progress in Brain Research. Elsevier; 2009 [cited 2023 Dec 11]. p. 339–51.Available from: https://www.sciencedirect.com/science/article/pii/S0079612309177233

4. Koch Fager S, Fried-Oken M, Jakobs T, Beukelman DR. New and emerging access technologies for adults with complex communication needs and severe motor impairments: State of the science. Augment Altern Commun Baltim Md 1985 2019;35(1):13–25.

5. Card NS, Wairagkar M, Iacobacci C, et al. An Accurate and Rapidly Calibrating Speech Neuroprosthesis. N Engl J Med 2024;391(7):609–18.

6. Willett FR, Avansino DT, Hochberg LR, Henderson JM, Shenoy KV. High-performance brain-to-text communication via handwriting. Nature 2021;593(7858):249–54.

7. Willett FR, Kunz EM, Fan C, et al. A high-performance speech neuroprosthesis. Nature 2023;620(7976):1031–6.

8. Littlejohn KT, Cho CJ, Liu JR, et al. A streaming brain-to-voice neuroprosthesis to restore naturalistic communication. Nat Neurosci 2025;28(4):902–12.

9. Pandarinath C, Nuyujukian P, Blabe CH, et al. High performance communication by people with paralysis using an intracortical brain-computer interface. eLife 2017;6:e18554.

10. Metzger SL, Littlejohn KT, Silva AB, et al. A high-performance neuroprosthesis for speech decoding and avatar control. Nature 2023;620(7976):1037–46.

11. Moses DA, Metzger SL, Liu JR, et al. Neuroprosthesis for Decoding Speech in a Paralyzed Person with Anarthria. N Engl J Med 2021;385(3):217–27.

12. Wairagkar M, Card NS, Singer-Clark T, et al. An instantaneous voice-synthesis neuroprosthesis. Nature 2025;1–8.

13. Jude JJ, Levi-Aharoni H, Acosta AJ, et al. An intuitive, bimanual, high-throughput QWERTY touch typing neuroprosthesis for people with tetraplegia [Internet]. 2025 [cited 2025 Jun 26];2025.04.01.25324990. Available from: https://www.medrxiv.org/content/10.1101/2025.04.01.25324990v1

14. Weiss JM, Gaunt RA, Franklin R, Boninger ML, Collinger JL. Demonstration of a portable intracortical brain-computer interface. Brain-Comput Interfaces 2019;6(4):106–17.

15. Bacher D, Jarosiewicz B, Masse NY, et al. Neural Point-and-Click Communication by a Person With Incomplete Locked-In Syndrome. Neurorehabil Neural Repair 2015;29(5):462–71.

16. Dekleva BM, Weiss JM, Boninger ML, Collinger JL. Generalizable cursor click decoding using grasp-related neural transients. J Neural Eng 2021;18(4):0460e9.

17. Brandman DM, Hosman T, Saab J, et al. Rapid calibration of an intracortical brain–computer interface for people with tetraplegia. J Neural Eng 2018;15(2):026007.

18. Singer-Clark T, Hou X, Card NS, et al. Speech motor cortex enables BCI cursor control and click. J Neural Eng 2025;22(3):036015.

19. Perge JA, Homer ML, Malik WQ, et al. Intra-day signal instabilities affect decoding performance in an intracortical neural interface system. J Neural Eng 2013;10(3):036004.

20. Downey JE, Schwed N, Chase SM, Schwartz AB, Collinger JL. Intracortical recording stability in human brain-computer interface users. J Neural Eng 2018;15(4):046016.

21. Sponheim C, Papadourakis V, Collinger JL, et al. Longevity and reliability of chronic unit recordings using the Utah, intracortical multi-electrode arrays. J Neural Eng 2021;18(6):066044.

22. Ali YH, Bodkin K, Rigotti-Thompson M, et al. BRAND: a platform for closed-loop experiments with deep network models. J Neural Eng 2024;21(2):026046.

23. Young D, Willett F, Memberg WD, et al. Signal processing methods for reducing artifacts in microelectrode brain recordings caused by functional electrical stimulation. J Neural Eng 2018;15(2):026014.

24. The CMU Pronouncing Dictionary [Internet]. [cited 2025 May 13];Available from: http://www.speech.cs.cmu.edu/cgi-bin/cmudict

25. Park J, Kim K. g2pe [Internet]. 2019;Available from: https://github.com/Kyubyong/g2p

26. Su J, Lu Y, Pan S, Murtadha A, Wen B, Liu Y. RoFormer: Enhanced Transformer with Rotary Position Embedding [Internet]. 2023 [cited 2025 May 13];Available from: http://arxiv.org/abs/2104.09864

27. Zhang B, Sennrich R. Root Mean Square Layer Normalization [Internet]. 2019 [cited 2025 May 13];Available from: http://arxiv.org/abs/1910.07467

28. Simeral JD, Kim S-P, Black MJ, Donoghue JP, Hochberg LR. Neural control of cursor trajectory and click by a human with tetraplegia 1000 days after implant of an intracortical microelectrode array. J Neural Eng 2011;8(2):025027.

29. Nuyujukian P, Fan JM, Kao JC, Ryu SI, Shenoy KV. A High-Performance Keyboard Neural Prosthesis Enabled by Task Optimization. IEEE Trans Biomed Eng 2015;62(1):21–9.

30. Hochberg LR, Serruya MD, Friehs GM, et al. Neuronal ensemble control of prosthetic devices by a human with tetraplegia. Nature 2006;442(7099):164–71.

31. Kao JC, Nuyujukian P, Ryu SI, Shenoy KV. A High-Performance Neural Prosthesis Incorporating Discrete State Selection With Hidden Markov Models. IEEE Trans Biomed Eng 2017;64(4):935–45.

32. Nuyujukian P, Kao JC, Fan JM, Stavisky SD, Ryu SI, Shenoy KV. Performance sustaining intracortical neural prostheses. J Neural Eng 2014;11(6):066003.

33. Gilja V, Pandarinath C, Blabe CH, et al. Clinical translation of a high-performance neural prosthesis. Nat Med 2015;21(10):1142–5.

34. Gilja V, Nuyujukian P, Chestek CA, et al. A high-performance neural prosthesis enabled by control algorithm design. Nat Neurosci 2012;15(12):1752–7.

35. Sussillo D, Stavisky SD, Kao JC, Ryu SI, Shenoy KV. Making brain–machine interfaces robust to future neural variability. Nat Commun 2016;7(1):13749.

36. Pyglet [Internet]. Available from: https://github.com/pyglet/pyglet

